# Time-restricted eating promotes sustained fat loss in *Drosophila*

**DOI:** 10.64898/2026.08.05.742897

**Authors:** Jared A. Gatto, Timothy Y. Chang, Wendy C. Kanmogne, Serena Pen, Letitia R. Bortey, Jessica N. Kwon, Lia Mahal, Holden S. Kim, Leah Berhanu, Fatma Oduk, Megan R. Rabon, Scarlet J. Park, Erin L. Barnhart, William W. Ja, Nicholas Stavropoulos, Julie C. Canman, Mimi Shirasu-Hiza

## Abstract

While current therapeutics restricting calorie intake, such as GLP-1 agonists, induce fat loss for many people, they are ineffective for others and concerns remain about their long-term effects on health, particularly loss of lean muscle mass. Moreover, many quickly regain fat if they stop treatment. In contrast, time-restricted eating does not restrict calorie intake but instead restricts the time window for eating and prevents obesity in mice and humans. Here we investigated the effects of intermittent <u>T</u>ime-<u>R</u>estricted <u>F</u>eeding (iTRF), which extends lifespan and delays markers of aging, on stored fat in *Drosophila*. Ten days of iTRF caused significant fat loss relative to *ad lib* diet, an effect that persisted even after return to *ad lib* diet. Unlike iTRF-induced lifespan extension, iTRF-induced fat loss did not depend on circadian-regulated autophagy. iTRF treated both diet-induced and genetically induced obesity and significantly reduced lipid droplet size in the fat body (adipose tissue). Instead of causing muscle loss, iTRF increased total and muscle-specific protein levels and enhanced flight performance, suggesting a shift in body composition. We found that iTRF evoked fasting-induced hyperactivity, partially mediated by octopamine, the fly ortholog of the human stress hormone norepinephrine. Ablation of octopaminergic neurons (OANs) prevented iTRF-mediated effects: fat loss, increased protein, and enhanced flight performance. Our results suggest that *Drosophila* iTRF causes rapid, permanent fat loss and increased muscle function through a “fight or flight” response. Understanding the mechanisms driving differences between *Drosophila* and human responses to TRE could be critical for identifying effective therapeutic targets for obesity.

## INTRODUCTION

Today, we live in a society with unprecedented access to food. Not only is the American diet extremely calorie-rich, but many Americans consume calories from the time they wake up until they go to sleep, and even in the middle of the night (1–3). These habits lead to excessive caloric intake and high rates of obesity, which have become a major clinical problem in the United States. Obesity rates in the United States have skyrocketed over the past 30 years, rising from 22.9% to 40.3% in adults aged 20 years or older (4) – similar changes have occurred in children (5). While the introduction of obesity-treating drugs, such as GLP-1 agonists, has led to a decline in rising rates of obesity (6), the long-term effects remain unknown because metabolic rebound occurs when these drugs are discontinued (7,8). In addition, the long-term effects of GLP-1 agonists on health and aging remain unclear. There are concerns that GLP-1 agonists can lead to loss of lean muscle mass and accelerate sarcopenia, or age-associated muscle decline (9,10). Obesity is associated with comorbidities such as type 2 diabetes and metabolic syndrome (11). While ongoing research focuses on the mechanisms by which dietary components affect obesity, via overconsumption of calories (12) or lifestyle (13,14), the mechanisms driving fat loss due to timed cycles of fasting remain unclear.

Recent studies have suggested that time-restricted eating or intermittent fasting diets, which restrict eating to specific time windows and alternate those with periods of fasting, can confer significant long-term health benefits. In mice, these benefits include reduced oxidative stress and inflammation, decreased insulin resistance, and lower blood sugar (15–18). Similar outcomes have been observed in humans on short-term time-restricted diets, including improved 24-hour glucose levels, reduced blood pressure, and weight loss (19–22). There is evidence that timing-based diets can also prevent metabolic disease. In mice and humans, studies have shown that restricting feeding to part of the active phase without changing caloric intake reduced fat levels, protected against a high-fat diet, and reversed obesity (15–18,20,22–24). It has been hypothesized that regular and consecutive feeding and fasting cycles induce shifts in lipid metabolism and energy homeostasis that drive weight loss and other health benefits.

While short-term clinical studies lasting three months or less have consistently shown health benefits (25–28), long-term results have been much less consistent (29,30). This inconsistency may partly result from poor patient adherence; patients find it difficult to follow time-restricted diets for more than three months (31). Some studies have attempted to identify modified conditions, like personal preferences for the eating window, that might still provide health benefits (28,32). The difference between the results for human patients and model organisms may simply reflect differences in compliance; alternatively, there may be inherent biological differences between humans and model organisms in their metabolic responses to consecutive periods of fasting and feeding. Identifying those differences and understanding the mechanisms underlying long-term physiological changes in model organisms could provide insights into potential therapeutic interventions for humans.

*Drosophila melanogaster* is a short-lived, genetically tractable model organism with high evolutionary conservation in most major metabolic pathways (33–35). More recently, time-restricted diets have been shown to confer multiple health benefits on *Drosophila*. Our lab developed a time-restricted diet called intermittent Time-Restricted Feeding (iTRF), which, when administered for 30 days between early adulthood and midlife, extends lifespan and delays aging compared with an *ad libitum* (AL) diet with constant access to food (36). For AL diets, to control for the stimulation of diet changes, flies were moved to new food vials at the same times of day as those on iTRF diet. In our previous publication (36), we found that lifespan extension and delayed aging require a functional circadian clock and autophagic components. Other labs have shown that different forms of TRF extend lifespan (37,38), ameliorate metabolic alterations on high-fat diets (39), promote muscle function in obesity models (40), decrease oxidative stress (41), and attenuate age-related cardiac decline (42). While these studies have established many TRF-induced health benefits in *Drosophila*, the direct effects of time-restricted diets on fat (triacylglyceride) storage and the regulatory pathways that drive changes in lipid metabolism have not been well characterized.

In this study, we examined changes in triacylglyceride (TAG) storage during and after 10 days of iTRF diet. Like mammals, *Drosophila* mainly store energy (fat) as triacylglyceride in lipid droplets (34). We found that flies previously treated with iTRF starved more quickly than those previously on an *ad lib* diet, indicating a rapid and significant metabolic change. Using thin-layer chromatography to directly measure TAG levels, we found that iTRF flies had much lower TAG levels than *ad lib* controls, even though iTRF flies ate more than *ad lib* controls, along with increased protein levels. iTRF did not appear to affect the kinetics of TAG usage, as iTRF and *ad lib* flies used TAGs at the same rate during starvation. iTRF-induced fat loss occurred because iTRF flies lost fat during the fasting period and did not restore their fat to *ad lib* levels during the refeeding periods. Unexpectedly, iTRF-induced fat loss appeared to be permanent. iTRF-induced fat loss occurred in unmated females, mated females, and mated males and, unlike iTRF-mediated lifespan extension (36), did not require circadian or autophagic components. iTRF-induced fat loss occurred in a genetic model of obesity and enabled both the prevention and treatment of diet-induced obesity. When we imaged lipid droplets in the abdominal fat body, the main adipose tissue, we found that iTRF flies did not have fewer lipid droplets relative to *ad lib* flies, but the size of individual lipid droplets was significantly reduced. We also found that iTRF increased muscle-specific proteins in the thorax and improved flight performance. We found that fasting during the day induced hyperactive behavior, which is known to be dependent on octopamine signaling. Octopamine is the fly analog for norepinephrine, the mammalian stress hormone. While fasting-induced hyperactivity (“timed exercise”) was not sufficient to induce fat loss, ablating octopamine-producing neurons (OANs) prevented iTRF-induced fat loss, protein gain, and improved flight performance, and dietary octopamine supplementation to OAN-ablated flies restored fat loss. Taken together, these results suggest that fasting-stimulated stress hormone octopamine is necessary for iTRF-induced fat loss and that OAN signaling mediates iTRF effects on physiology.

## RESULTS

### Flies lose fat during intermittent time-restricted feeding and do not regain it

To examine how intermittent Time-Restricted Feeding (iTRF) influences metabolism in *Drosophila*, we compared flies on iTRF or *ad lib* diets. We used age-matched, mated female flies for our experiments unless otherwise stated. In brief, 10-day-old iTRF flies were fasted for 20 hours and then fed for 28 hours; for most experiments, this cycle was repeated five times (**Fig. 1A**). *Ad lib* (AL) flies had continuous access to food. As we previously reported, iTRF flies compensated for the fast period during refeeding by consuming more over 48 hours than their AL counterparts in cycles 2 and 3 of iTRF (36). Here, using the capillary feeder (CaFe) assay, we confirmed that, while iTRF flies ate the same amount as AL flies in Cycle 1, they consumed significantly more than AL flies in Cycles 4 and 5 (**Fig. 1B**). To assess overall metabolic changes caused by iTRF, we compared AL and iTRF flies for their ability to survive starvation after five cycles of fasting and feeding. We found that iTRF flies died significantly faster from starvation than AL flies (**Fig. 1C**), suggesting that some aspects of their metabolism and energy storage are affected. This also indicated that five cycles of fasting and feeding are sufficient to observe metabolic phenotypes.

**Figure 1.**
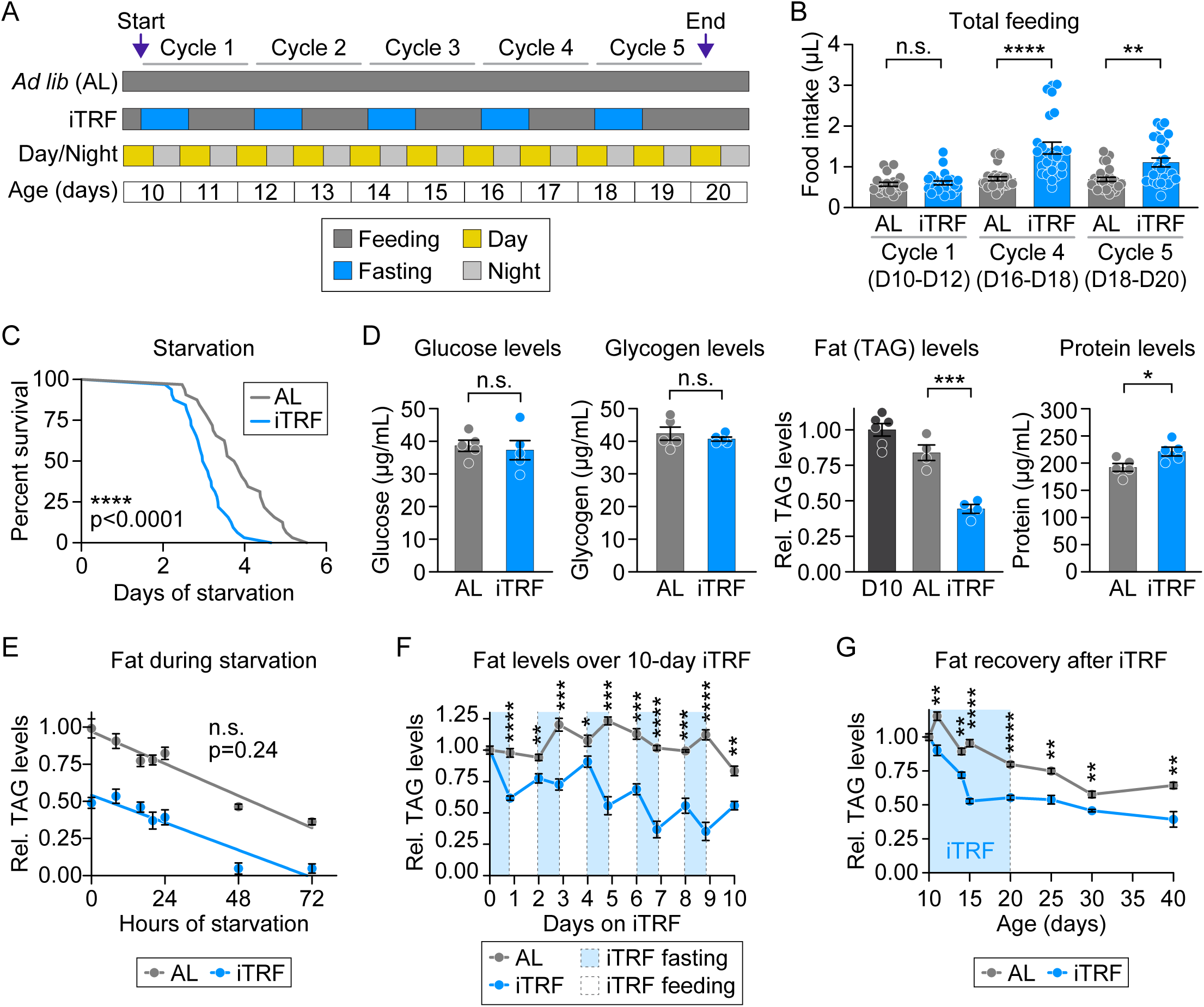
Flies lose fat during intermittent, time-restricted feeding and do not regain it. **(A)** Schematic of intermittent time-restricted feeding (iTRF) protocol. At 10 days old, flies are divided into ad libitum (AL, gray) and iTRF (blue) groups. iTRF flies fast for 20 hours and feed for 28 hours in each 48-hour cycle; *ad lib* flies have continuous access to food but are switched to new vials of the same food when iTRF flies are switched to different foods. Typical 10-day iTRF protocol has 5 fasting/feeding cycles. **(B)** iTRF flies ate the same amount as *ad lib* flies (gray) in cycle 1 (p>0.05) and ate significantly more in cycles 4 (p<0.0001) and 5 (p<0.01). Feeding was measured via Activity Recording Capillary Feeder (ARC) assay. Each point = 1 fly; n = 20-28 flies per condition. **(C)** iTRF flies died faster from starvation than AL controls (p<0.0001); n = 32 flies per condition. **(D)** While glucose and glycogen levels were unchanged after 10 days of iTRF (p>0.05), iTRF flies had lower fat levels (p<0.001) and higher protein levels (p<0.05). n = 10 flies per replicate per condition; 4-5 replicates per condition. **(E)** AL and iTRF flies lost fat at the same rate during starvation (p=0.2422). n = 10 flies per replicate per condition; 4 replicates per condition; each point = average across replicates. **(F)** iTRF flies lost fat during fasting phase (light blue) and did not fully regain fat during refeeding phase (white) (p<0.05 at all time points after 0). n = 10 flies per replicate per condition; 4 replicates per condition; each point = average across replicates. **(G)** Even after switched back to *ad lib* diet, iTRF flies never completely regain fat stores relative to *ad lib* diet (p<0.01 for all time points after 0); n = 10 flies per replicate per condition; 4 replicates per condition; each point = average across replicates. For p-values on graphs, n.s.: >0.05; *: ≤0.05. **: ≤0.01; ***: ≤0.001; ****: ≤0.0001. p-values were obtained by two-tailed independent t-test (B, D, F, G), logrank analysis (C), and simple linear regression (E); each experiment was performed 2-3 times.

To characterize metabolic pathways that might be impacted by iTRF, we examined three common energy substrates in *Drosophila*—glycogen, protein, and fat—along with glucose levels and body mass measurements, comparing AL and iTRF flies after five cycles of the dietary regimen. As with humans, the major form of fat storage in flies is triacylglyceride (TAGs), which becomes depleted after starvation (43). Using colorimetric assays to measure glucose, glycogen, and protein levels and thin-layer chromatography (TLCs) to measure fat levels (triacylglycerides or TAGs) (**Fig. S1A**), we found that female flies treated with either AL and iTRF diets did not exhibit differences in body mass (**Fig. S1B**), glucose (**Fig. 1D**), or glycogen (**Fig.1D**). In contrast, iTRF-treated females had significantly lower fat levels and significantly higher protein levels than those on AL diet (**Fig. 1D**). Males on iTRF diet lost fat but did not gain protein **(Fig. S1C, S1H)**. Thus, for both males and females, even though iTRF flies ate more than AL flies overall, the repeated cycles of fasting and feeding led to significant fat loss. Unexpectedly, female flies also gained protein. This result suggested that these repeated cycles of fasting and feeding led to a significant change in fat utilization (catabolism) or storage (anabolism), consistent with findings in other model organisms (15,16,44).

To test whether, after five cycles of fasting and feeding, iTRF flies burned their fat stores at a faster rate than ad lib flies, we subjected AL and iTRF flies to starvation conditions and measured fat loss over time. We found that flies that had previously been on either AL or iTRF diet lost fat at identical rates during starvation assays (**Fig.1E**), suggesting that iTRF does not enhance the kinetics of fat catabolism. To test if iTRF flies were altered in fat storage, we measured fat levels during the 10-day iTRF period. We observed that iTRF flies lost fat during each fast period and, although they ate more than AL flies during the post-fast feeding period, did not replenish their fat stores to AL levels (**Fig. 1F**).

To determine the time needed for iTRF flies to replenish their fat stores to AL levels or higher (as seen with human “yo-yo dieting”) (45), we transferred iTRF flies to AL diets after five cycles of fasting and feeding and measured their fat levels for the next 20 days. Unexpectedly, we found that iTRF flies never replenished their fat stores to the same levels observed in AL flies (**Fig. 1G**), a pattern not observed with humans on TRE diets (29). This result indicates that iTRF-induced fat loss in *Drosophila* is long-lasting and suggests that the underlying mechanism(s) continue to limit fat storage in *Drosophila* even after iTRF has been discontinued.

### iTRF fat loss is not specific to sex or reproductive status

We have typically focused on mated females for iTRF experiments. Because metabolism can differ with reproductive status (mated or virgin females) (46) and between sexes (male or female) (47,48), we set out to test whether the physiological effects of 5 cycles of iTRF, including lifespan and fat loss, were specific to mated females. We confirmed that, as previously reported (49), virgin females had a longer lifespan than mated females; we also found that virgin females experienced additional lifespan extension with 30 days of iTRF compared with virgin females on an AL diet (**Fig. S1D**). Virgin females further exhibited decreases in starvation survival time (**Fig. S1E**) and fat loss (**Fig. S1F**) after 10 days of iTRF, relative to virgin females on AL diet. Consistent with results from mated and virgin females, iTRF-treated mated males also starved more quickly (**Fig. S1G**) and had significantly lower fat levels relative to AL males (**Fig. S1H**). Therefore, the metabolic changes induced by iTRF are general and not specific to mating status or sex.

### iTRF-induced fat loss does not require circadian or autophagy mechanisms

In a previous study, we showed that autophagy and circadian clock components are required for iTRF-induced lifespan extension (36). Here, we used the same genetic manipulations, RNAi-mediated knockdown of autophagy component *atg1* and loss-of-function mutation of circadian component period (*per^01^* mutant), to test if these autophagy and circadian clock components were also required for iTRF-induced fat loss. We found that all control genotypes and experimental flies with knockdown of *atg1* starved to death more quickly (**Fig. 2A**) and had lower fat levels (**Fig. 2B**) after 10 days of iTRF, relative to those on AL diet. While iTRF-treated *per^01^* mutants did not starve more quickly, they still had significant fat loss relative to *per^01^* mutants on AL diet (**Fig. 2C**, **2D**). These results indicate that, unlike iTRF-induced lifespan extension, autophagy and circadian components are not required for iTRF-induced fat loss. Unexpectedly, this result also suggests that differences in starvation time do not always reflect differences in fat levels; *per^01^* mutants lost fat on iTRF, but this fat loss did not accelerate death by starvation.

**Figure 2.**
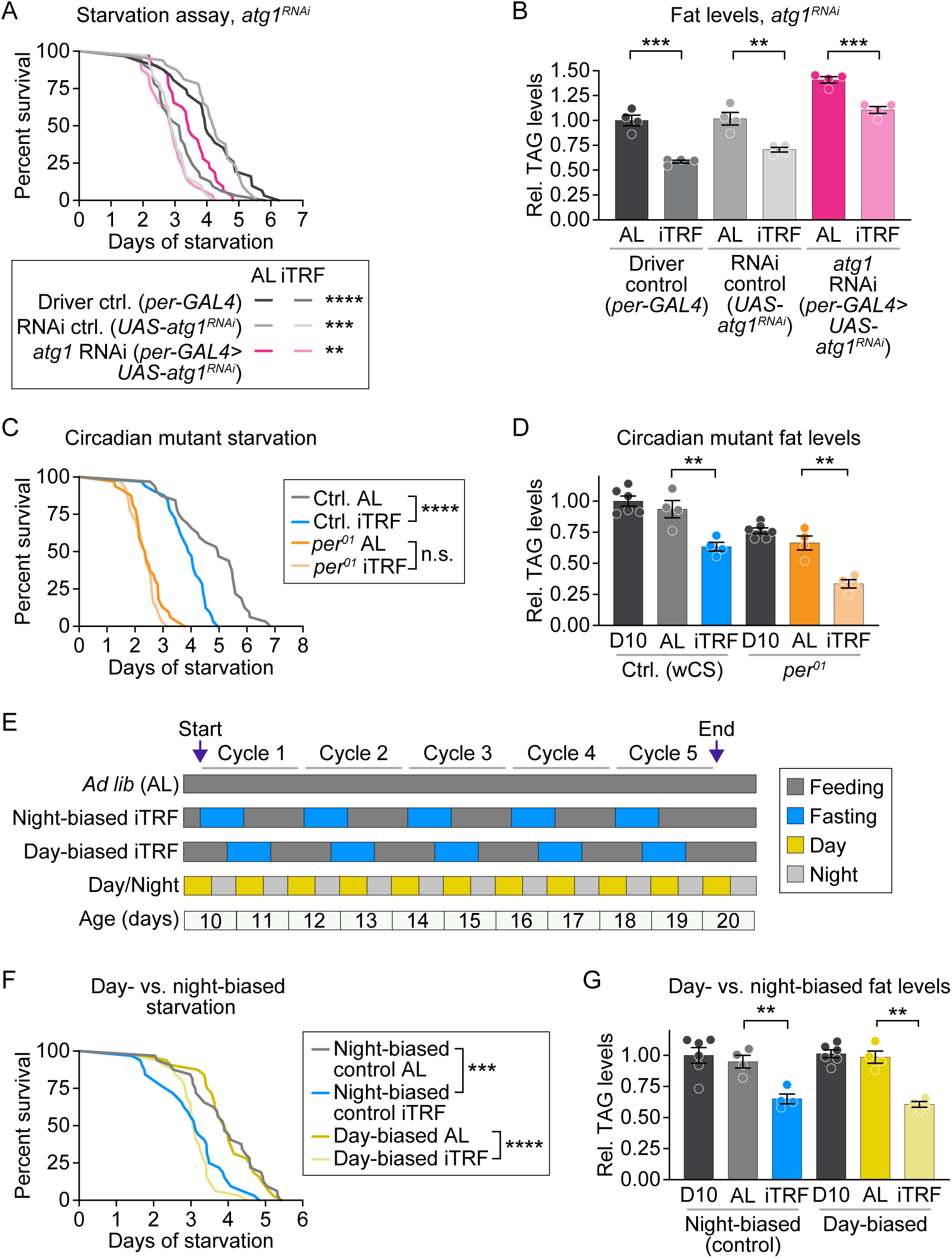
iTRF-induced fat loss does not require essential circadian or autophagy components. **(A)** RNAi of *atg1* did not inhibit iTRF-induced starvation sensitivity. Both genetic controls (gray) and flies with circadian knockdown of *atg1* (pink) died more quickly from starvation if previously treated with iTRF (light gray, light pink) relative to *ad lib* diet (dark gray, dark pink); p<0.0001 for *per*-*GAL4* controls; p<0.001 for UAS-*atg1*^RNAi^ controls; p<0.01 for *per*-*GAL4>*UAS-*atg1*^RNA^ KD flies); n = 32 flies per condition per genotype. **(B)** RNAi of *atg1* did not inhibit iTRF-induced fat loss. Both genetic controls (gray) and flies with circadian knockdown of *atg1* (pink) lost fat with iTRF treatment relative to *ad lib* diet (p<0.001 for *per*-*GAL4* controls and *per*-*GAL4>*UAS-*atg1*^RNA^ KD; p<0.01 for UAS-*atg1*^RNAi^ controls). n = 10 flies per replicate per condition per genotype; 4 replicates per condition. (**C)** Arrhythmic *per^01^* mutants treated with iTRF (light orange) did not die faster during starvation than those treated with *ad lib* diet (dark orange, p>0.05), unlike genetic controls (iTRF, blue; *ad lib,* gray; p<0.0001); n=32 flies/condition/genotype. **(D)** Arrhythmic *per^01^* mutants lost fat on iTRF (light orange) relative to *ad lib* diet (dark orange, p<0.01), similar to genetic controls (iTRF, blue; *ad lib*, dark gray; p<0.01). n=10 flies/replicate/condition/genotype; 4 replicates/condition. **(E)** Schematic of day-biased vs. night-biased iTRF and *ad lib* diets. **(F)** Flies treated with day-biased iTRF (light yellow) died faster during starvation than day-biased *ad lib* flies (dark yellow, p<0.0001), similar to flies treated with night-biased iTRF (blue) relative to night-biased *ad lib* flies (dark gray, p<0.001); n=32 flies/condition. **(G)** Day-biased flies lost fat on iTRF (light yellow) relative to ad lib diet (dark yellow, p<0.01), similar to night-biased controls (iTRF, blue; *ad lib*, gray; p<0.01). n = 10 flies per replicate per condition; 4 replicates per condition. For p-values on graphs, n.s.: >0.05; *: ≤0.05. **: ≤0.01; ***: ≤0.001; ****: ≤0.0001. p-values were obtained by two-tailed independent t-test (B, D, G) and logrank analysis (A, C, F); each experiment was performed 3 times.

We further investigated circadian requirements by shifting the fasting period to center around the day, which is the fly’s active period. Previously, we showed that, when the fasting period is centered around the day (“day-biased”) rather than the night (“night-biased”), iTRF did not extend lifespan (36). To confirm that iTRF-induced fat loss did not depend on circadian time of day, we measured fat levels after four dietary treatments: night-biased iTRF; night-biased AL diet; day-biased iTRF; and day-biased AL diet (**Fig. 2E**). We found that day-biased iTRF flies, like night-biased iTRF flies, starved more quickly (**Fig. 2F**) and lost more fat (**Fig. 2G**) than their matching AL controls. Taken together, these data indicate that the mechanisms responsible for iTRF-induced fat loss do not require circadian or autophagy components and are therefore distinct from the mechanisms driving iTRF-induced lifespan extension.

### iTRF stimulated fat loss in models of both genetic obesity and diet-induced obesity

Proposed health benefits of fasting-based diets include weight loss and the prevention of obesity (50). To test if iTRF could promote *Drosophila* fat loss even under obesogenic conditions, we tested both genetic and diet-induced models of obesity. To genetically induce high levels of stored fat, we used the well-characterized *brummer* (*bmm*) mutant, which has elevated triacylglyceride levels compared with its isogenic control (48,51,52). *bmm* encodes a lipase, orthologous to mammalian ATGL, which is important for cytosolic lipolysis. This process breaks down stored triacylglycerides, releasing free fatty acids that serve as substrates for mitochondrial beta-oxidation (48).

We hypothesized that, if iTRF-induced fat loss depended on cytosolic lipolysis and Bmm lipase, *bmm*^01^ mutants would not exhibit iTRF-induced sensitivity to starvation and would be resistant to iTRF-induced fat loss. To test this, we subjected both *bmm*^01^ and its control (*bmm*^rev^) to 10 days of either iTRF or AL diet, then measured starvation survival time and fat levels. Consistent with previous studies and their obese state (48,52), after 10 days of either iTRF or AL diet, *bmm^01^* mutants had longer starvation survival time (**Fig. 3A**) and higher TAG levels (**Fig. 3B**) compared to their wild-type isogenic controls. When we compared iTRF-treated *bmm*^01^ mutants to AL-treated *bmm*^01^ mutants, we found that iTRF treatment was still able to induce shorter starvation survival time (**Fig. 3A**) and fat loss (**Fig. 3B**) relative to AL diet. These results, showing fat loss in this genetic model of obesity, indicate that *brummer* is not required for iTRF-induced metabolic effects.

**Figure 3.**
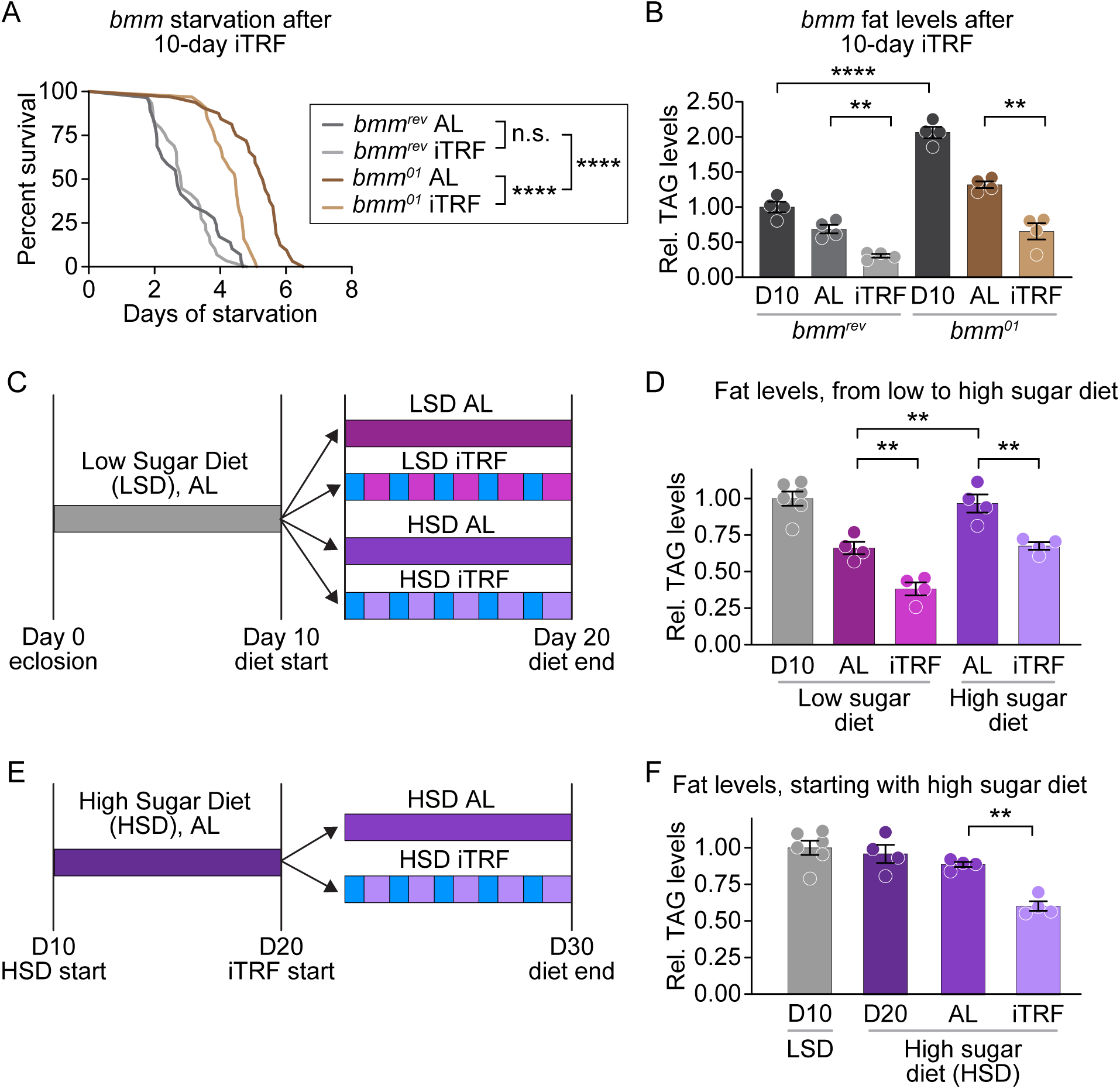
iTRF stimulated fat loss in models of both genetic obesity and diet-induced obesity. **(A)** *brummer* mutants (brown) were resistant to starvation relative to genetic controls (gray, p<0.0001) and died faster when previously treated with iTRF (light brown) than *ad lib* diet (dark brown, p<0.0001), unlike genetic controls (light gray vs. dark gray, p>0.05); n = 29-32 flies per condition per genotype. **(B)** *brummer* mutants had more fat than controls on day 10 (dark gray vs. very dark gray, p<0.0001) but still lost fat on iTRF (light brown vs. dark brown, p<0.01), similar to controls (light gray vs. dark gray, p<0.01). n = 10 flies per replicate per condition per genotype; 4 replicates per condition. **(C)** Schematic of dietary conditions for flies fed a low-sugar diet (LSD) until 10 days old and split into 4 cohorts: AL LSD, iTRF LSD, AL high-sugar diet (HSD), and iTRF HSD. **(D)** Whether on iTRF or *ad lib* diets, HSD flies (purple) had higher fat levels on day 20 relative to LSD flies (pink vs. purple, p<0.01); both LSD and HSD flies lost fat on iTRF relative to *ad lib* diets (light vs. dark hues, p<0.01). n = 10 flies per replicate per condition; 4-6 replicates per condition. **(E)** Schematic showing dietary switch performed at day 20; flies were given AL HSD from day 10 to day 20 and split into AL HSD and iTRF HSD groups for day 20 to day 30. **(F)** Flies on iTRF HSD (light purple) had lower fat levels relative to AL HSD (dark purple, p<0.01). n = 10 flies per replicate per condition; 4-6 replicates per condition. For p-values on graphs, n.s.: >0.05; *: ≤0.05. **: ≤0.01; ***: ≤0.001; ****: ≤0.0001. p-values were obtained by logrank analysis (A) and two-tailed independent t-test (B, D, F); each experiment was performed at least 2 times.

Similar to mammals on obesogenic Western diets, *Drosophila* fed a high-sugar diet (HSD, 0.7 M sucrose) have high fat levels, increased insulin resistance, and decreased fertility (34,53). We first set out to test if iTRF prevented this diet-induced increase in TAG levels. Flies raised on a low-sugar diet (LSD) for ten days were treated with four different diets over the next 10 days: low sugar diet (LSD) AL or iTRF, or high sugar diet (HSD) AL or iTRF (**Fig. 3C**). As expected, flies on low-sugar diet had significantly lower TAG levels on iTRF than AL diet (**Fig. 3D**). While flies on high-sugar AL diet had higher TAG levels than flies on a low-sugar AL diet, flies on a high-sugar iTRF diet had the same TAG levels as flies on low sugar AL diet, which were lower than those on high-sugar AL diet (**Fig. 3D**). This result suggests that iTRF can prevent the increase in fat stores due to high sugar diet.

Although not consistent across all reports, earlier studies with mice have indicated that time-restricted feeding can sometimes be less effective at treating obesity than at preventing it (54,55). To test if iTRF can induce fat loss in Drosophila that are already obese, we raised flies on high-sugar AL diet for 10 days to increase TAG levels and then treated these flies with either high-sugar AL diet or high-sugar iTRF for 10 more days (**Fig. 3E**). We found that flies on high-sugar iTRF diet had significantly decreased TAG levels relative to those on high-sugar AL diet, even though they started iTRF with already elevated TAG levels (**Fig. 3F**). Taken together, these results suggest that iTRF can not only prevent the increase in TAG levels due to a high-sugar diet but also decrease high TAG levels that have already resulted from a high-sugar diet.

### iTRF-treated flies have smaller lipid droplets in their fat body cells and increased muscle proteins in their thoraces

Each segment of the fly (the head, thorax, and abdomen) contains measurable fat and protein; the adult *Drosophila* adipose tissue or fat body is distributed throughout the body (56), and each tissue contains protein in various forms, such as eggs in the abdomen or indirect flight muscles in the thorax (57). To determine whether there is segment specificity for iTRF-induced fat loss and iTRF-induced protein gain, we dissected flies after 10 days of iTRF or AL diet and measured TAG and protein levels for each segment (head, thorax, and abdomen) (**Fig. 4A**). All segments lost fat during iTRF, with the largest fat decrease in the abdomen, which contains more fat body tissue than any other segment (56) (**Fig. 4B**); in addition, thoraces gained considerable protein during iTRF, which contains major protein reservoirs in the indirect flight muscles (58) (**Fig. 4C**). These results suggest that iTRF induced changes in fat dynamics within the abdominal fat body, and iTRF induced changes in protein dynamics within the thorax.

**Figure 4.**
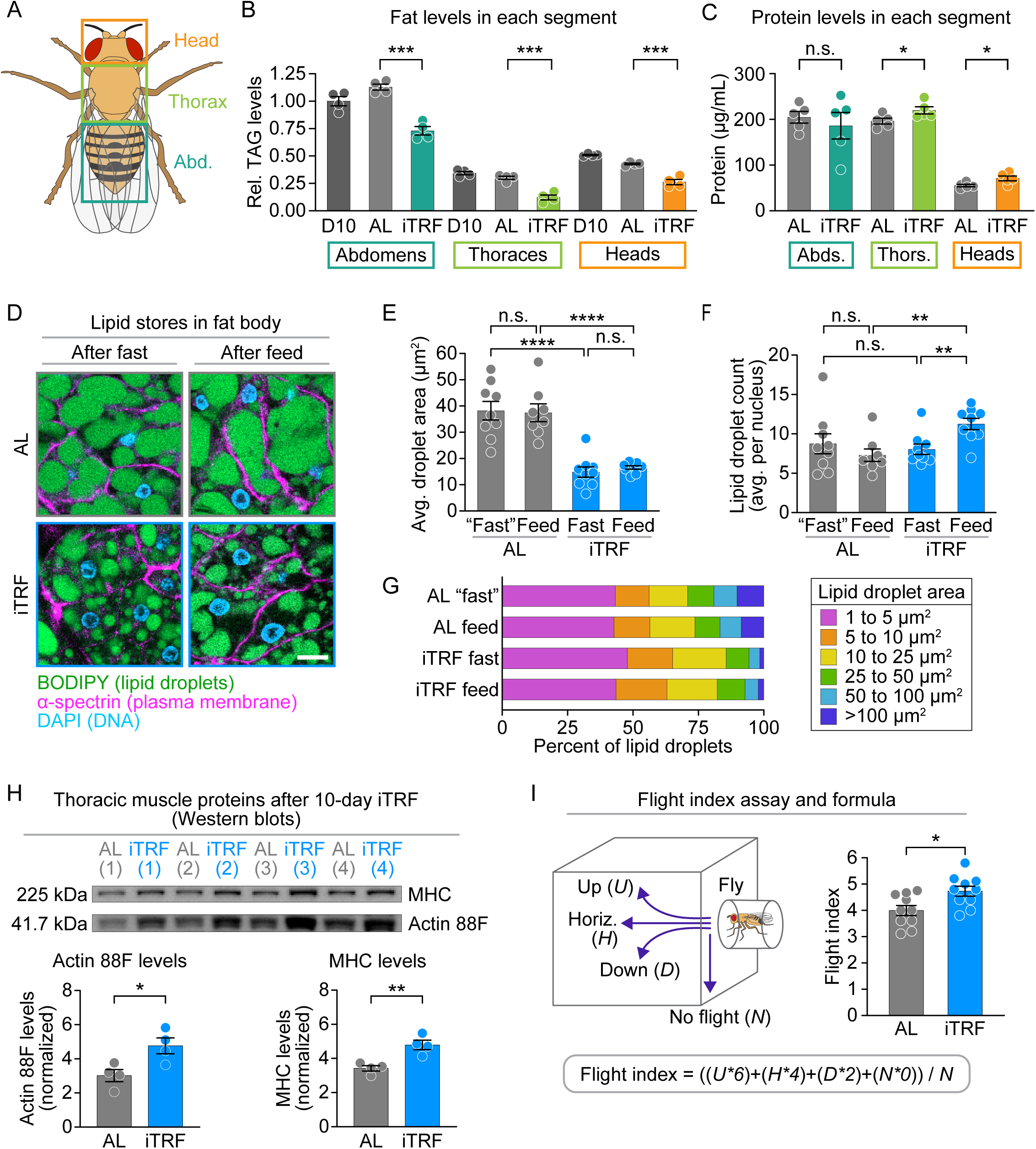
iTRF-treated flies have smaller lipid droplets in their fat body cells and increased muscle proteins in their thoraces. **(A)** Schematic of different body segments tested for fat levels comparing *ad lib* (AL) and iTRF diets, showing head (orange), thorax (green), and abdomen (teal). **(B)** For each body segment, iTRF flies (blue) had lower fat levels relative to *ad lib* flies (gray, p<0.001). n = 40 heads, 15 thoraces, or 10 abdomens per replicate per condition; 4 replicates per condition for each segment. **(C)** Thoraces and heads from iTRF flies (blue) had higher levels of protein than those from *ad lib* flies (gray, p<0.05); there was no difference in abdominal segments (p>0.05). n = 40 heads, 30 thoraces, or 20 abdomens per replicate per condition; 5 replicates per condition for each segment. **(D)** Representative regions of interest of AL and iTRF fat body sections after the last fasting cycle (after fast) and feeding cycle (after feed); green = BODIPY stain (fat), blue = DAPI stain (nuclei), magenta = ɑ-spectrin (membrane). **(E)** ITRF flies (blue) had smaller lipid droplets relative to AL flies (gray) in both fast and feed phases (p<0.0001 for both); there was no difference within diet conditions (p>0.05). Each point = average of 6 ROIs per fat body per condition; 8-9 replicates per condition. **(F)** While the number of lipid droplets per nucleus was not different between iTRF and AL flies at the end of fasting (p>0.05), iTRF flies had more lipid droplets per nucleus at the end of feeding phase than AL flies (p<0.01). Each point = average of 6 ROIs per image per condition; 8-9 replicates per condition. **(G)** Histogram analysis of lipid droplets binned by size (area) demonstrates more small lipid droplets (pink) and fewer very large lipid droplets (dark blue) for iTRF flies relative to ad lib flies. **(H)** Quantification of western blot of myosin heavy chain (MHC) and actin 88F shows that dissected thoraces from iTRF flies (blue) had increased levels of actin 88F (left, p<0.05) and MHC (right, p<0.01) relative to those from AL flies (gray). n=30 thoraces per replicate per condition; 5 replicates per condition; values normalized to Ponceau values in individual lanes. **(I)** Schematic for flight index assay, along with the formula used to calculate flight index based on released flies’ initial flight direction. iTRF flies (blue) had better flight performance than ad lib flies (gray, p<0.05). n = 10 flies per replicate per condition; 10 replicates per condition. For p-values on graphs, n.s.: >0.05; *: ≤0.05. **: ≤0.01; ***: ≤0.001; ****: ≤0.0001. p-values were obtained by two-tailed independent t-test (B, C, E, F, H, I); each experiment was performed at least 2 times.

The *Drosophila* fat body performs many functions similar to those of the mammalian liver, including glycogen storage, nutrient sensing, and detoxification, and also acts as the main adipose tissue, storing fat as triglycerides (35,59,60). To analyze changes in fat storage during iTRF, we dissected fat bodies from flies after 10 days of either AL or iTRF diets. We collected samples after the final iTRF fasting and refeeding (**Fig. 4D**) periods. We used a lipophilic dye (BODIPY) to stain individual lipid droplets, in parallel with staining of the plasma membrane (anti-α-spectrin) and DNA (DAPI), with regions of interest (ROIs) chosen through an imaging process and thresholding pipeline **(Fig. S2A, S2B)**. We found that, both at the end of the fasting period and at the end of the feeding period, iTRF flies have smaller lipid droplets than AL flies (**Fig. 4E**). We also found that, while the number of lipid droplets per nucleus was similar between iTRF and AL flies at the end of the fasting period, at the end of the next feeding period, iTRF flies had more lipid droplets per nucleus than AL flies (**Fig. 4F**). Size histogram analysis demonstrated that, at this time point, there were many more very small lipid droplets than in AL or feeding iTRF samples (**Fig. 4G**). Taken together, these results suggest that iTRF reduces the size of lipid droplets in the fat body relative to AL diet and that fasting depletes very small lipid droplets first, while refeeding restores those very small lipid droplets.

To address potential tissue-specific effects of iTRF-induced protein gain, we next investigated the thorax. The thorax had significantly increased protein levels during iTRF (**Fig. 4C**) and is a tissue enriched for muscle cells that power movement and flight (58), suggesting that increases in thoracic protein might reflect higher levels of muscle-specific proteins. To test whether muscle-specific proteins increased during iTRF, we used western blot analysis to measure levels of myosin heavy chain (MHC) and actin 88F, the predominant actin isoform in thoracic indirect flight muscles. In Drosophila, indirect flight muscles (IDMs) rely on MHC and actin 88F to contract, deform the thorax, and power high-frequency wingbeats (58,61). After 10 days of iTRF, we dissected thoracic samples for western blotting and found that both actin 88F and MHC protein levels were significantly increased in iTRF compared to their AL controls (**Fig. 4H**), suggesting that iTRF could promote muscle cellular function.

To test whether this increase in muscle-specific protein translates to better functional performance, we used the flight index assay, a well-described behavioral experiment in which flies are released into an arena and flight directionality is recorded; it has been previously used in flies under TRF conditions (40,62). A weighted scoring formula is used to indicate flight performance (**Fig. 4I**). After 10 days of iTRF, we subjected both AL and iTRF flies to a flight index assay and found that iTRF flies performed better than their AL controls (**Fig. 4I**), suggesting that iTRF improves muscle performance by increasing muscle-specific proteins in the thorax.

### iTRF-induced hyperactivity is not sufficient for fat loss

Given that 10 days of iTRF increases muscle-specific proteins (**Fig. 4G**) and flight performance (**Fig. 4H**), we tested if iTRF increases locomotor activity. To measure locomotor activity, we placed individual flies into Drosophila Activity Monitors (DAMs), which continuously measure activity throughout the day and night (63), and treated them with either iTRF or AL diet (**Fig. 5A**). We compared locomotor activity over five cycles of iTRF (ten days), binning activity into four categories based on iTRF diet and time of day: daytime fasting, nighttime fasting, daytime feeding, nighttime feeding. We found that iTRF significantly increased average activity, especially during daytime fasting and nighttime feeding (**Fig. 5A**). During daytime fasting, all iTRF cycles showed a significant increase in activity (**Fig. 5A**), consistent with starvation-induced hyperactivity, as previously reported (64,65). While this hyperactivity was significantly decreased as soon as the lights turned off (nighttime), increased locomotor activity was also observed during cycle 5 nighttime fasting, as well as both daytime feeding (cycles 2, 3, 5) and nighttime feeding (all cycles) (**Fig. 5A**). These results demonstrate that iTRF significantly increased locomotor activity relative to AL diet.

**Figure 5.**
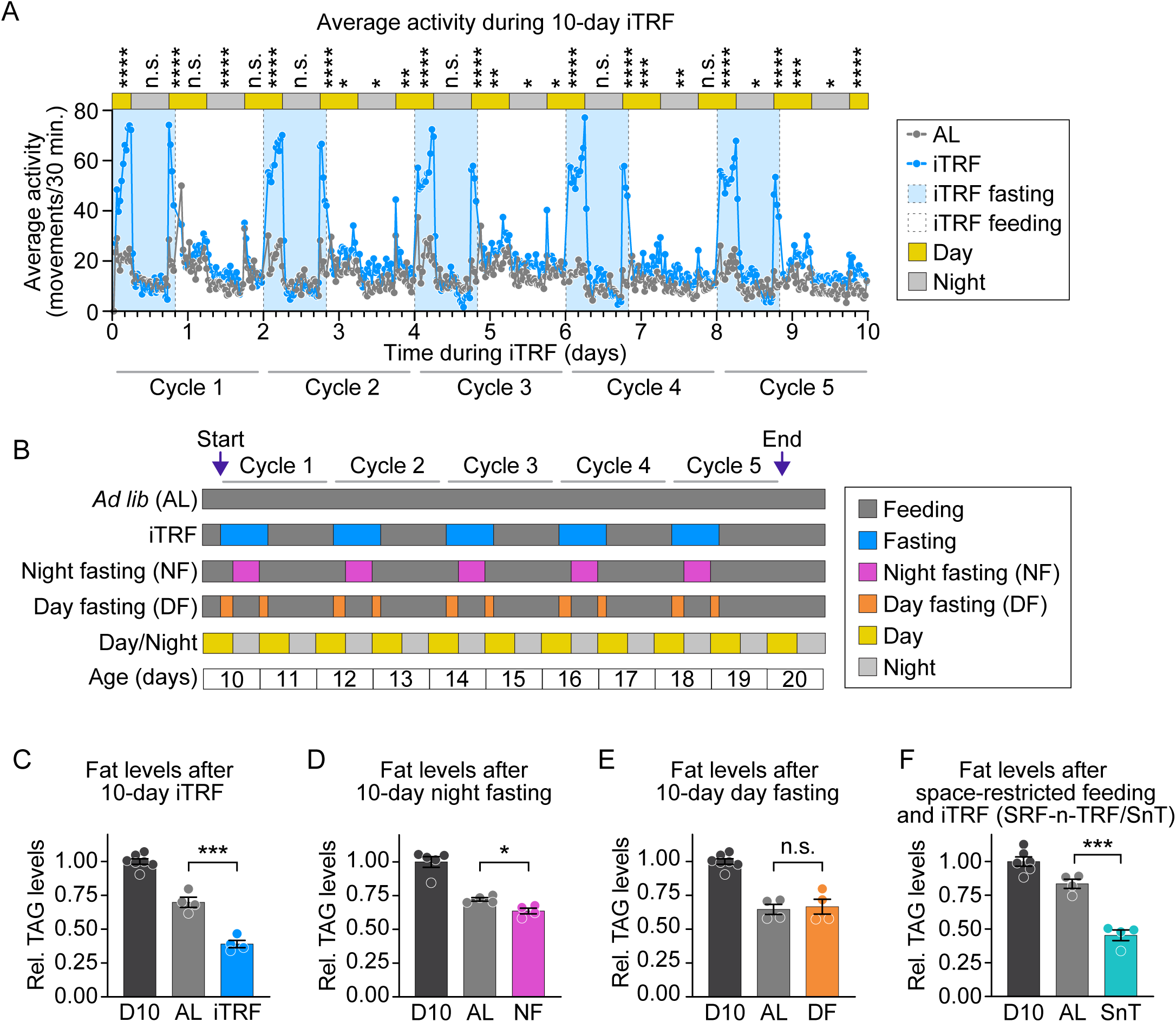
iTRF-induced hyperactivity is not sufficient for fat loss. **(A)** iTRF increased activity, particularly during day fasting and night feeding periods (see p-values corresponding to times of day). Average activity via DAM activity measurement was plotted over the 10-day iTRF period for AL (gray) and iTRF (blue) flies, shown with iTRF fasting periods (light blue box), feeding periods (white box), day (yellow bar), and night (gray bar). Dashed lines represent time of transfer for both AL and iTRF flies to new DAM tubes. p-values were obtained by binning activity per condition, time of day, and fasting/feeding periods. n = 32 flies per replicate per condition; 3 replicates per condition. **(B)** Schematic of conditions to test the effect of activity on fat loss by separating iTRF into night fasting condition (NF, low activity) and day fasting condition (DF, hyperactivity). Relative to AL flies (gray): **(C)** iTRF flies (blue) had lower fat levels (p<0.001); **(D)** night-fasted flies (NF, pink) had lower fat levels (p<0.05); and **(E)** day fasted flies (DF, orange) did not have lower fat levels (p>0.05). **(F)** Restricting movement by restricting space did not affect iTRF-induced fat loss; flies undergoing space-restricted feeding and time-restricted feeding (SRFnTRF) still had lower fat levels than AL flies (p<0.001). For these experiments (**C-F**), n = 10 flies per replicate per condition; 4 replicates per condition. For p-values on graphs, n.s.: >0.05; *: ≤0.05. **: ≤0.01; ***: ≤0.001; ****: ≤0.0001. p-values were obtained by two-tailed independent t-tests unless otherwise indicated. Each experiment was performed at least 2 times.

Because of this iTRF-induced increase in locomotor activity, we investigated whether iTRF also altered sleep patterns. Sleep and activity levels are typically, but not always, inversely related; moreover, changes in sleep can affect metabolism in both model organisms and humans (66). In line with standards in the field of *Drosophila* sleep, we defined sleep as a cessation of activity lasting 5 minutes or longer (67,68). We found that iTRF flies experienced less average sleep, lower sleep bout number, and lower sleep bout length than AL flies during day fasting (**Fig. S3A-C**). Other conditions (nighttime fasting, daytime feeding, and nighttime feeding) showed different effects. During night fasting, flies had significant sleep loss in 3 out of 5 cycles (**Fig. S3A**), while also exhibiting reduced sleep bout number (**Fig. S3B**) and increased sleep bout length (**Fig. S3C**), indicating more consolidated sleep – this effect decreased toward the end of the 10-day period. Day and night feeding showed similar trends: decreased average sleep (**Fig. S3A**), fewer sleep bouts (**Fig. S3B**), but longer sleep bouts (**Fig. S3C**) toward the beginning of iTRF, which diminishes toward the end. Overall, this suggests that the spike in activity levels during daytime fasting has a physiological effect on iTRF flies.

Given that increased activity (exercise) can lead to fat loss in both model organisms and humans (69), we hypothesized that iTRF-induced hyperactivity causes fat loss. Other time-restricted diets have also been reported to increase locomotor activity (41,42). To test this, we split iTRF treatment into its day and night components: flies treated with a control AL diet, the iTRF diet, daytime fasting (associated with starvation-induced hyperactivity), or nighttime fasting (not associated with starvation-induced hyperactivity) (**Fig. 5B**). We first confirmed that iTRF treatment of flies in the narrow DAM tubes still caused significant fat loss (**Fig. S3D**). Using standard vials, we found that, while iTRF (**Fig. 5C**) and nighttime fasting (**Fig. 5D**) significantly reduced fat levels, daytime fasting (with high activity levels alone) did not (**Fig. 5E**). This result suggests that hyperactivity alone was not sufficient to cause fat loss. We also tested whether immobilizing the flies prevented fat loss during iTRF. Using space-restricted conditions to limit locomotor movement as previously described (70) for both iTRF and AL flies, we observed that space-restricted feeding and time-restricted feeding (SRF-n-TRF) flies still lost fat relative to space-restricted AL flies (**Fig. 5F**). These findings suggest that the iTRF-induced increase in locomotor activity does not drive iTRF-induced fat loss.

### OAN-mediated octopamine secretion is required for iTRF-induced fat loss

We next set out to inhibit hyperactivity using genetic ablation of cells that regulate this behavioral response. During stress such as starvation, Akh, or adipokinetic hormone (orthologous to mammalian glucagon), is released by adipokinetic hormone-producing cells (APCs) (71). Akh signals to the fat body to catabolize fat stores for energy and is a key upstream activator of *brummer* (71,72). Akh also signals to octopaminergic neurons (OANs) to produce octopamine, or OA (orthologous to mammalian norepinephrine), a neurohormone that mediates starvation-induced stress responses like increased hyperactivity and inhibition of egg laying (73) (**Fig. 6A**). Since the Akh-OA pathway is crucial to the starvation response, we asked if ablating APCs or OANs would affect iTRF-induced fat loss.

**Figure 6.**
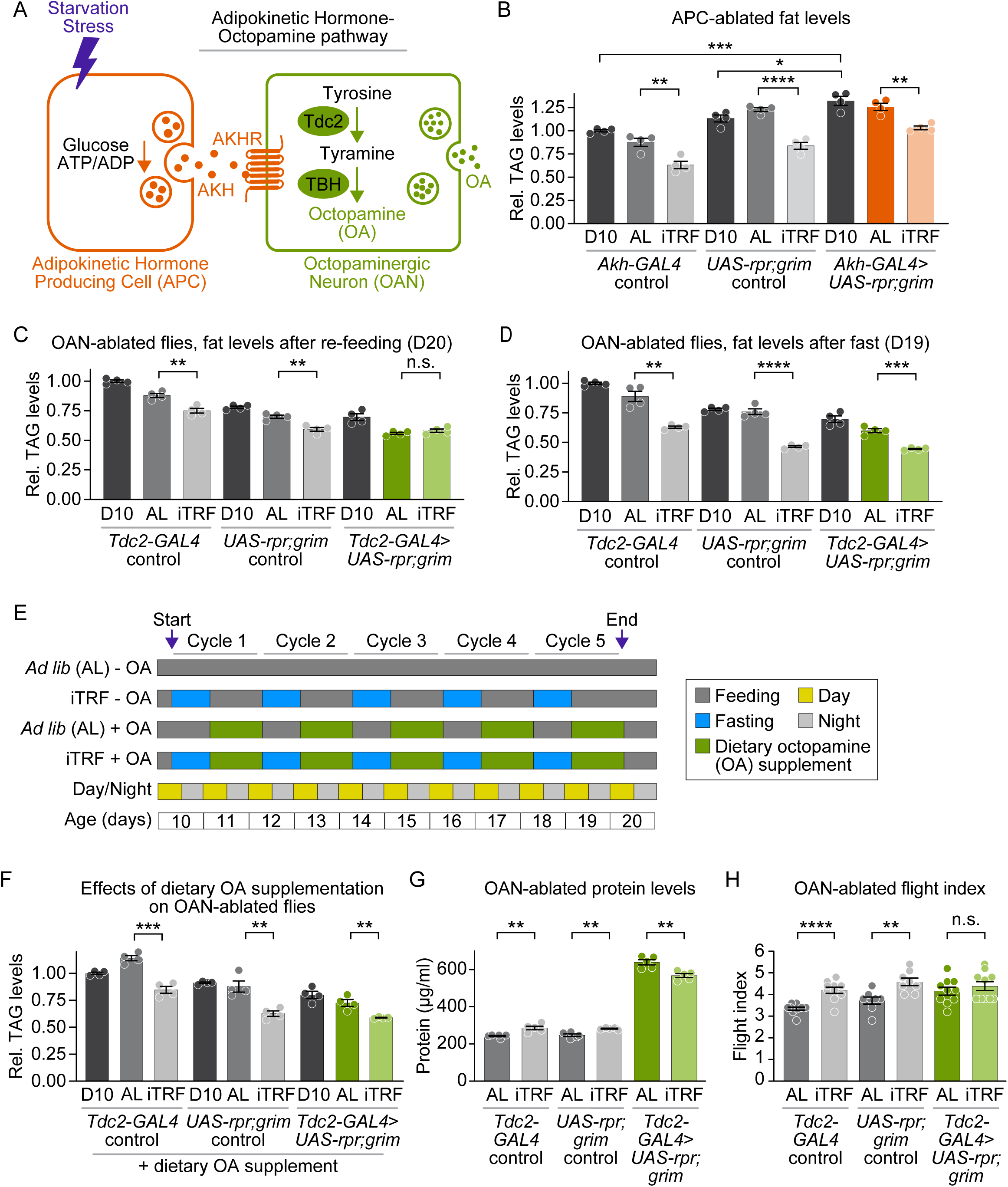
OAN-mediated octopamine secretion is required for iTRF-induced fat loss. **(A)** Schematic showing the AKH-OA signaling pathway, important for regulating energy homeostasis during starvation. **(B)** Flies with ablated AKH-producing cells (APCs, orange) had more fat than genetic controls at day 10 (dark grays; p<0.001 for *Akh-GAL4* control; p<0.05 for *UAS-rpr; grim* control) and had lower fat levels on iTRF vs. AL diets (light orange vs. dark orange, p<0.01), similar to genetic controls (light grays vs. dark grays, p<0.01 for *Akh-GAL4* control, p<0.0001 for *UAS-rpr; grim* control). **(C)** In contrast with genetic controls, which had lower fat levels on iTRF compared to AL diet (light grays vs. dark grays, p<0.01 for both *Tdc2-GAL4* and *UAS-rpr; grim* controls), flies with ablated octopaminergic neurons (OANs, *Tdc2-GAL4>UAS-rpr; grim*) did not have lower fat levels after iTRF (light green) compared to AL diet (dark green, p>0.05). **(D)** OAN-ablated flies still lost fat during the iTRF fasting period (dark green vs. light green, p<0.01). **(E)** Schematic showing dietary supplementation of octopamine during feeding periods with non-supplemented controls. **(F)** Unlike non-supplemented controls (C), OAN-ablated flies (*Tdc2-GAL4> UAS-rpr; grim*) supplemented with dietary octopamine had lower fat levels after iTRF (light green) compared to AL diet (dark green, 0.001), similar to genetic controls (iTRF vs. AL, p<0.001 for *Tdc2-GAL4* control, p<0.01 for *UAS-rpr; grim* control). **(G)** While controls had higher protein levels on iTRF compared to AL diets (p<0.01 for both controls), OAN-ablated flies had lower protein levels on iTRF compared to AL diet (light green vs. dark green, p<0.01). **(H)** While genetic controls had improved flight performance on iTRF (light grays) compared to AL diet (dark grays, p<0.0001 for *Tdc2-GAL4* control, p<0.01 for *UAS-rpr; grim* control), OAN-ablated flies did not (light green vs. dark green, p<0.05 for *Tdc2-GAL4*>*UAS-rpr; grim*). n = 10 flies per replicate per condition per genotype (B, C, D, F, G, H); 4 replicates per condition per genotype (B, C, D, F); 5 replicates per condition per genotype (G); 8-10 replicates per condition per genotype (H). For p-values on graphs, n.s.: >0.05; *: ≤0.05. **: ≤0.01; ***: ≤0.001; ****: ≤0.0001. p-values were obtained by two-tailed independent t-tests. Each experiment was performed at least 2 times.

To ablate Akh-Producing Cells (APCs) or Octopaminergic Neurons (OANs), we drove two pro-apoptotic transgenes (UAS-*reaper* and UAS-*grim*) with either *Akh*-GAL4 driver (for APCs) or *Tdc2*-GAL4 driver (for OANs), as previously reported (73). To confirm ablation, we used *Drosophila* activity monitors to measure the locomotor activity of 3-day old flies in starved and fed conditions, comparing ablated flies to genetic controls. We plotted their average activity for 4 consecutive days **(Fig. S4A, S4B)** and took aggregate measurements of activity during day and night periods over a 48-hour period **(Fig. S4C, S4D)**. Ablating APCs is known to suppress starvation-induced hyperactivity (74). Consistent with this, we found that, while their genetic controls displayed starvation-induced hyperactivity, APC-ablated flies did not **(Fig. S4A, S4C)**. The effect of OAN ablation using pro-apoptotic constructs on overall activity has not been previously described, though mutants defective in octopamine production do not undergo starvation-induced hyperactivity (75). We found that, while OAN-ablated flies had a reduced starvation response compared to their controls (**Fig. S4C, S4D)**, they still experienced starvation-induced hyperactivity **(Fig. S4B, S4D)**. As validation of the ablation, we note that OAN-ablated flies exhibited a well-known phenotype for loss of octopamine: egg-laying deficiency and large abdomens filled with unlaid eggs **(Fig. S4E)** (76).

We next measured TAG levels in control flies and those with ablated APCs or ablated OANs after 10 days of AL or iTRF. We found that both control flies and flies with ablated APCs still lost fat compared to flies on an AL diet (**Fig. 6B**). These results confirmed that another genetic obesity model could undergo iTRF-induced fat loss and provided further evidence that starvation-induced hyperactivity is not responsible for iTRF-induced fat loss. In contrast, while all genetic controls for OAN ablation lost fat on iTRF compared to AL diet, we found that flies with ablated OANs did not lose fat on iTRF (**Fig. 6C**). Males with ablated OANs did lose fat during iTRF **(Fig. S4F)**, suggesting a sex-specific difference in OAN ablation and iTRF-induced fat loss. Overall, this suggests that either octopamine or tyramine, two neurohormones produced by the OANs, is necessary for female flies to lose fat on iTRF.

To test if flies with ablated OANs have the same fat levels on iTRF and AL diets because they were unable to lose fat during the fasting period or unable to regain it during refeeding, we measured fat levels after the final fasting period, before the final refeeding period. We found that flies with ablated OANs did indeed lose fat during fasting with iTRF, similar to controls (**Fig. 6D**). This result suggested that OAN-ablated flies lost fat during fasting but, in contrast to controls, fully replenished their fat stores upon refeeding.

Because OANs produce other neurohormones such as tyramine (73) and because this phenotype could be due to a developmental effect, we tested whether dietary octopamine supplementation of OAN-ablated flies could induce fat loss during iTRF. We measured TAG levels for OAN-ablated flies and their controls on iTRF or AL diets, with vehicle alone or dietary octopamine supplementation (**Fig. 6E**). We found that dietary octopamine supplementation was sufficient to induce fat loss in OAN-ablated flies on iTRF, similar to controls (**Fig. 6F**). This result suggests that OAN-mediated octopamine secretion is both necessary and sufficient for iTRF-induced fat loss for OAN-ablated flies.

Given the effect of OAN-ablation on fat loss, we next tested if OAN-ablation altered other iTRF-mediated effects, such as increased protein levels and improved flight performance. As a positive control, we subjected APC-ablated flies and their controls to both protein measurements and flight index assays after 10-days of iTRF and found that APC-ablated flies, along with their controls, exhibited increased protein levels **(Fig. S4G)** and improved flight performance **(Fig. S4H)**. In contrast, while their controls gained protein and improved flight performance after iTRF, OAN-ablated flies lost protein after iTRF (**Fig. 6G**) and iTRF did not improve flight performance compared to AL (**Fig. 6H**). This suggests that OAN is also implicated in other iTRF-mediated effects beyond fat loss, indicating that OANs play a major role in mediating iTRF’s effects on physiology.

## DISCUSSION

While the link between intermittent fasting and metabolism has been observed previously (18,23,39,40,44,50), this study directly investigates the effects of iTRF on fat levels in *Drosophila*. We found that, after 10 days on iTRF diet, flies died faster from starvation than flies on AL diet, indicating possible changes in metabolism. We showed that iTRF flies, regardless of sex or reproductive status, had significantly lower triacylglyceride (TAG) levels after the feeding periods of iTRF. TAG levels in iTRF mated female flies were not fully restored to AL levels after refeeding, and fat loss persists for at least 30 days even after returning to an AL diet. The mechanisms driving iTRF-mediated fat loss appeared to differ from those driving iTRF-mediated lifespan extension (36), which include circadian and autophagic processes. We also found that fly models of both genetic and diet-induced obesity lost fat after 10 days of iTRF diet, indicating a potential therapeutic approach. These changing lipid dynamics could be observed in the *Drosophila* fat body. Lipid droplets in iTRF-treated flies were significantly smaller than those in *ad lib* flies, both during the fasting and refeeding periods. While iTRF and *ad lib* flies had the same number of lipid droplets per nucleus at the end of the fasting period, iTRF flies had more lipid droplets, particularly very tiny lipid droplets, by the end of the feeding period relative to ad lib flies or fasting iTF flies. Finally, we discovered that when neurons responsible for octopamine signaling are ablated, flies do not lose fat on iTRF – this can be restored with octopamine supplementation. This result suggests that octopamine, a stress-induced hormone that plays a role similar to that of norepinephrine in mammals, is both required and sufficient for iTRF-induced fat loss.

A key finding of this study is that iTRF-induced fat loss appeared to be irreversible. Strikingly, fat stores were not replenished even long after the end of iTRF treatment. These results contrast with previous research in humans and other models, such as mice (77,78). In human trials, all measures of time-restricted eating (TRE)-driven fat loss—such as body fat percentage, body mass, and visceral fat—return to baseline within three months, suggesting that ongoing TRE is necessary to maintain benefits (22,79). Although *Drosophila* fat bodies are orthologs of mammalian adipose tissue, differences in their structure and function may explain this difference in response to TRE. Unlike mammalian adipose tissue, the *Drosophila* fat body is a versatile metabolic center that coordinates fat, liver, and immune functions and undergoes rapid structural changes in response to fasting and nutrient states (35,59,60). As seen in iTRF, these include structural differences in lipid droplet size and number per cell. Consistent with these observations, iTRF not only produced persistent reductions in fat stores but also increased whole-body and muscle-specific protein content, suggesting that iTRF induced broader remodeling of body composition rather than simply depleting lipid reserves. Future research examining how iTRF influences fat body organization, tissue architecture, and protein accumulation will shed light on the mechanisms that sustain this shift in body composition and fat loss.

In *Drosophila*, the enzymes tyrosine decarboxylase 2 (Tdc2) and tyramine β-hydroxylase (Tbh) work together in specific neurons to convert tyrosine into tyramine and octopamine, vital neurohormones that act systemically to regulate locomotion, feeding behavior, and reproductive physiology (76,80). Octopamine serves as the functional *Drosophila* analog of norepinephrine (NE), coordinating arousal, metabolism, and energy utilization across tissues in response to physiological demand (73,80). Octopamine is present only at trace levels in humans and does not play a major signaling role in human physiology, highlighting a potential limitation in translating our work in *Drosophila* to humans. Nevertheless, NE is integral to the sympathetic tone in humans, controlling fat catabolism (81). Our work highlights the potential of investigating NE and its impact on TRE-induced benefits in mammalian models. Since octopamine controls many physiological processes in *Drosophila*, it will be important to determine which function(s) contribute to iTRF-induced fat loss and in which tissues it’s required.

Using a *Drosophila* model of iTRF-induced fat loss offers unique advantages, including the ability to track metabolic parameters and overall physiology in a short-lived, genetically tractable model. This approach provides a clear advantage for future studies with high statistical power. Our model can help elucidate the underpinnings of the effects of time-restricted feeding on metabolism, reproduction, and health span. Future work on potential therapeutic targets against obesity will aid the fight against the obesity epidemic and could offer insight to other research groups using time-restricted feeding in a similar fashion.

## MATERIALS AND METHODS

### Fly Strains

*w1118 Canton-S* (*wCS*) flies were used as our “wild-type” strain throughout this paper. *UAS*-*atg1*-RNAi (#44034), *Akh-GAL4* (#25683), and *Tdc2-GAL4* (#9313) were obtained from the Bloomington Stock Center. *period* (*per*^01^) mutants and *period*-*GAL4* lines were obtained from J. Giebultowicz. *bmm*^01^ and *bmm*^rev^ lines were obtained from E. Rideout, with permission from R. Kühnlein. The UAS-*reaper*, UAS-*hid* double-construct line was obtained from W. Grueber. Controls for crosses include construct flies crossed with our *wCS* “wild type” strain.

### Fly Media

Developmental media were prepared using standard yeast-cornmeal-agar (Archon Scientific; glucose recipe: 7.6% glucose, 3.8% yeast, 5.3% cornmeal, 0.6% agar, 0.5% propionic acid, 0.1% methyl paraben, and 0.3% ethanol). Adult flies that eclosed within 24 hours were collected and transferred to ‘adult medium’ containing 4% dextrose, 2% sucrose, 5% cornmeal, 1% agar, and 3% yeast extract (Difco), supplemented with 1.5% methyl paraben mix (10% methyl paraben dissolved in ethanol) and 0.75% propionic acid. This was our standard experimental diet for AL and iTRF experiments. For our diet-induced obesity experiments, we used a control low-sugar diet with 1% agar, 8% brewer’s yeast, 2% yeast extract, 2% peptone, 5.13% sucrose (0.15M), 0.2% MgSO₄·7H₂O, 0.34% CaCl₂·2H₂O, 0.6% propionic acid, and 1% methyl paraben mix (10% methyl paraben dissolved in ethanol). The high-sugar diet contained the same ingredients and concentrations as the low-sugar control, except that sucrose was increased to 23.9% (0.7M). All percentages are weight/volume unless noted otherwise; methyl paraben mix, propionic acid, magnesium sulfate heptahydrate, and calcium chloride dihydrate are given as volume/volume. Fasting media consisted of 1% agar in ddH₂O and was prepared fresh daily. For octopamine supplementation, octopamine hydrochloride (Millipore Sigma, 00250) was added to the ‘adult medium’ at a final concentration of 4 mg/ml, as previously used.

### ARC Assay

Feeding data from individual flies was collected as previously described (82). Feeding medium consisted of a solution of 4% dextrose, 2% sucrose, and 3% yeast extract in ddH_2_O, filtered with a 0.2-μm cellulose acetate sterile syringe filter (VWR). Flies were transferred to fresh standard medium every other day until the start of iTRF. When the flies were 9-11 days old, the animals were loaded by mouth pipette into standard ARC chambers and allowed to acclimate overnight (approximately 18 hours) with access to the test diet in a glass capillary pipette. For later time points (four or five cycles of fasting), flies were placed into standard ARC chambers after completing three fasting cycles to begin feeding measurements. The capillaries were replaced daily with those containing fresh food or ddH2O. The meniscus level of each capillary was tracked throughout the entire feeding period at 1-second intervals. Drops in meniscus position above a pre-calibrated threshold were considered feeding events and feeding bouts less than 2 minutes apart were considered part of the same meal. The volume consumed during each feeding event was automatically calculated and compiled using a custom Python code.

### Starvation Assay

Starvation assays were performed as previously described (83). Newly eclosed flies (∼24 hours) were collected and allowed to mate for 48 hours. Female and male flies were separated, and females were maintained at a density of 25-30 flies per vial in a humidified, temperature-controlled (25 °C) incubator with a 12-hour light-dark (LD) cycle. About 10 days post-eclosion, flies were divided into AL and iTRF groups. After another 10 days on their respective diets, both groups were transferred to *Drosophila* activity monitor (DAM) tubes (5 mm) containing 1% agar and placed into a DAM system that tracks fly movement via infrared beams. Flies were monitored using a DAM2 system (Trikinetics) for 5-7 days. Activity data were summed into 15-minute increments, and survival curves were generated; time of death was recorded immediately after the last recorded movement. For each starvation experiment, 28-32 flies were used per group, and at least 3 independent replicates were conducted with different biological cohorts.

### Carbohydrate Measurements

Carbohydrate measurements were completed as previously described (83). To prepare samples for carbohydrate (glucose, glycogen) measurement, AL and iTRF flies were collected after 10 days on either dietary regimen in groups of 10 flies per replicate (5 replicates per condition/timepoint). Flies were washed several times in PBS, then homogenized in 200 µL PBS. Samples were centrifuged for 1 min at 4°C, and the supernatant removed and heated for 10 min at 70°C. Samples were then centrifuged again for 3 min at 4°C, and the supernatant removed and diluted 1:4 with PBS. Glycogen was measured via glucose detection with Glucose (HSK) Assay kit (Sigma GAHSK20) after 1 hour of 37°C amyloglucosidase digestion. Glucose values are from the non-digested samples from glycogen samples. Absorbance was measured using a plate reader at 25°C. Sample concentrations were determined using a series of 1:2 serial dilutions of glucose at known concentrations.

### Protein Measurements

Flies were collected after 10 days of iTRF and flash frozen. Flies were collected as follows: 10 bodies/replicate/condition, 40 heads/replicate/condition, 30 thoraces/replicate/condition, and 20 abdomens/replicate/condition. 5 replicates/condition were used for statistical analysis. Protein levels were measured with the Dilution-Free Rapid Gold BCA Protein Assay Kit (Pierce). Whole flies and body segments were homogenized in cold PBS, then centrifuged at 12,700 RPM at 4°C for 3 minutes. The supernatant was collected and diluted 1:15 to fall within the linear range of the BSA standard curve. 10 µL of standards and samples were pipetted onto a 96-well microplate (Costar), combined with the Pierce working reagent, and incubated at room temperature for 5 minutes. Absorbance was measured at 480 nm using an Infinite M Nano Plate Reader (Tecan). Sample absorbance values were corrected by subtracting the standard absorbance. A standard curve was generated from the standard measurements and used to determine the accurate protein concentrations of the samples.

### Mass and Triglyceride (Fat) Measurements

AL and iTRF flies were collected at various time points during the 10-day iTRF period in groups of 10 females per biological replicate (4 per condition), or 20 males per biological replicate (4 per condition), then weighed. Samples were homogenized in a 2:1 chloroform:ethanol solution to solubilize triglycerides, then centrifuged for 15 minutes at 4°C. Subsequently, samples were diluted 1:8 to fall within the linear range of our standards. They were then spotted onto a TLC plate (Sigma-Aldrich, Silica gel on TLC-PET foil 99577) and separated using a solvent mixture of 70 mL heptane, 30 mL diethyl ether, and 1 mL acetic acid. To normalize samples across plates, we used a “female fly homogenate” spotted in triplicate on all plates. This “female fly homogenate” was prepared from 10-day-old *wCS* flies (pre-iTRF), with 10 females per sample and 50 replicate samples. Extracted lipids across all samples were pooled, and serial dilutions were performed to determine a linear range. To visualize bands, plates were stained with 0.2% amido black (Napthol Blue Black, Sigma Product N-3393) in 1 M NaCl, dried overnight, and then imaged using an iBright CL 1500 system in the colorimetric western blot setting. Images were quantified using FIJI. Sample values were normalized to plate background, the “female fly homogenate” on their respective plates, and D10 (pre-iTRF) fat levels either within genotype or for controls to calculate relative triglyceride (TAG) levels for each sample.

### Flight index assay

Flight index assays were performed as previously described (40,62). Briefly, AL and iTRF flies after 10-days of iTRF (10 flies per vial, 8 vials/condition) are released at the center of a Plexiglas box with a light source positioned at the top (**Fig. 4H**). Initial flight direction per fly is scored and assigned a value: up (6), horizontal (4), down (2), or no flight (0). Flight indices are calculated by dividing the sum of the individual FI values by the number of individuals for each group.

### DAM activity analysis

Flies were collected 2 days post-eclosion from stock bottles and allowed to mate for 2 days in vials on our normal lab diet. Female flies were separated and allowed to age for an additional 6 days. At 10 days post-eclosion, 32 flies/condition/replicate, with 3 replicates, were loaded into Trikinetics DAM5 activity monitors in 5 mm plastic tubes and placed in a 25°C incubator. AL and iTRF were then unplugged from the DAM5 monitoring system and flipped together at fasting/feeding transitions. Flipping occurred within 30-45 minutes outside the incubator to minimize the period during which activity was not recorded. Fresh food and agar were made every fasting day, and fresh food was made every feeding day to ensure the food was hydrated. When the 10-day iTRF concluded, activity was divided into phases by time of day and feeding status (daytime fasting, nighttime fasting, daytime feeding, nighttime feeding). Activity counts were binned into 30-minute intervals by DAM file scan and then averaged across all replicates at each time point to create average activity for all flies.

### Sleep analysis

Sleep was measured in the same group of flies used for the DAM activity analysis. Beam break data were divided into 1-minute intervals with DAM File Scan and then processed using custom R code to automatically identify sleep, created by Dr. Nicholas Stavropolous. Sleep was defined as at least 5 minutes of inactivity.

### Virgin female lifespan analysis

*Drosophila* were reared from embryos in low-density bottles using standard yeast-cornmeal-agar media listed above (Archon Scientific). Virgin females were collected over a 1-2 day period post-eclosion and grouped with 25-30 females on adult medium containing 3% yeast extract. Age-matched control (mated) females were collected as usual and given 48 hours to mate, after which females and males were separated and placed on adult medium in groups of 25-30 flies. Both groups were kept in a humidified (65%) and temperature-controlled (25°C) incubator with a 12-hour light-dark cycle. All flies were raised under *ad lib* conditions until day 10 post-eclosion, when they were shifted to either *an ad lib* or an iTRF diet.

For consistency, AL control flies were transferred to fresh food at the same time as the experimental diet flies were switched to fasting media, and again when fasting ended, and flies were returned to regular adult media. Death was recorded at the time of transfer, and lifespan was analyzed using log-rank tests.

### Fluorescence staining of the fat body

AL and iTRF flies were collected and dissected at different time points during the 10-day iTRF period. Adult fat bodies were dissected in PBS and fixed in 4% PFA for 20 minutes. For plasma membrane staining, samples were blocked for 90 minutes in 5% natural goat serum (NGS) and incubated in anti-ɑ spectrin (3A9, DSHB, 3ug/mL) overnight at 4°C. Then, we used Invitrogen secondary anti-mouse IgG 594 (1:250). For lipid droplet visualization, we stained with BODIPY 505/515 (25ng/mL) in PBS for 30 minutes, then counterstained with 0.1% DAPI in PBST for 5 minutes x 3 times at room temperature. Stained fat bodies were then mounted in Slowfade^TM^ Gold Antifade (Invitrogen, S36937). Images were acquired with a Zeiss LSM-800 at 40X Magnification, using a 0.5 NA objective and standard laser lines (405, 488, 561, and 640 nm), with identical power settings for all samples. A minimum of 7 fat bodies per condition were used for each quantification, and at least three independent trials were performed for each experiment.

### Fat body image quantification

For quantification, a representative stack was chosen from each image – AL and iTRF fat bodies were max projected at the same depth in FIJI. 6 regions of interest (ROIs) were chosen in an unbiased manner **(Fig. S2A, S2B)** using the ɑ-spectrin (membrane) and DAPI (nucleus) channels. Each ROI was split into separate channels, and structures were quantified: lipid droplets (BODIPY) and nuclei (DAPI). Lipid droplet size and nuclei counts were collected per ROI and then averaged for each image.

### Statistical analysis and replicability

Statistical analysis was performed using GraphPad Prism 10. Significance is presented as *P* values (NS=*P*>0.05, \**P*<0.05, \*\**P*<0.01, \*\*\**P*<0.001, \*\*\*\**P*<0.0001). For comparison of two groups, we used unpaired, two-tailed *t*-tests when data met the criteria for parametric analysis (normal distribution and similar variance). All plotted values represented means, with error bars representing s.e.m. All biochemical experiments were performed with a minimum of 4 biological replicates, repeated in two to three independent trials. For comparison of survival curves (lifespan and starvation), log-rank (Mantel-Cox) analysis was used. Data from all trials are available in the supplemental Excel file.

## AUTHOR CONTRIBUTIONS

JAG, TYC, and MSH conceived and designed the experiments. Authors who contributed to the following experiments are listed in parentheses: husbandry (JAG, TYC, WCK, SP, LRB, JNK), iTRF and ad lib diets (JAG, TYC, WCK, SP, LRB, JNK), glucose assays (JAG, LM, MRR), glycogen assays (JAG, LM, MRR), thin-layer chromatography assays for fat (JAG, WCK, LRB, LM), BCA assays for protein (JAG, WCK, LM), starvation assays (JAG, TYC, LRB, LM), high sugar diet assays (JAG, WCK), fat and protein analysis per body segment (JAG, WCK, LRB), fat body dissections (TYC, SP, LRB), confocal immunofluorescence microscopy (TYC, SP), quantification of confocal immunofluorescence microscopy images (JAG), Western blots (JAG, TYC, WCK, HSK), flight index assays (JAG, TYC, WCK, JNK, LB, FO), iTRF in *Drosophila* activity monitors (JAG, TYC, HSK), timed exercise and space-restricted feeding (JAG, TYC, HSK), and genetic ablation experiments (JAG, TYC, JNK). SJP and WWJ performed feeding assays. NS wrote Matlab code and performed sleep analysis of iTRF DAM results. JAG, TYC, ELB, JCC, and MSH made intellectual contributions, designed the figures, and wrote the manuscript.

## ACKNOWLEDGMENTS

We thank all members of the Canman and Shirasu-Hiza labs for support, discussions, and feedback. We also thank Zola Stevens and Isaac Tom for their help with iTRF DAM assays and a very fun summer. Support for this research came from these sources: NIH F31AG079601 (JAG); NIH T32GM141882 (JAG, LM, MRR); NSF GRFP (TYC); Columbia University SURF Program (WCK, HSK); Barnard Summer Research Institute (JK); Columbia SPURS program, NIH R25NS076445 (LB); NIH R01NS121179 (ELB); NIH R01DC020031 (WWJ); NIH R01NS112844 (NS); NIH R35GM127049 (MSH); and NIH R01AG045842 (MSH).

## DATA AVAILABILITY

The authors declare that all data supporting the findings of this study are available, including replicate experiments, and will be made available upon reasonable request to the corresponding author, Mimi Shirasu-Hiza.

---

**Supplementary Figure 1.**
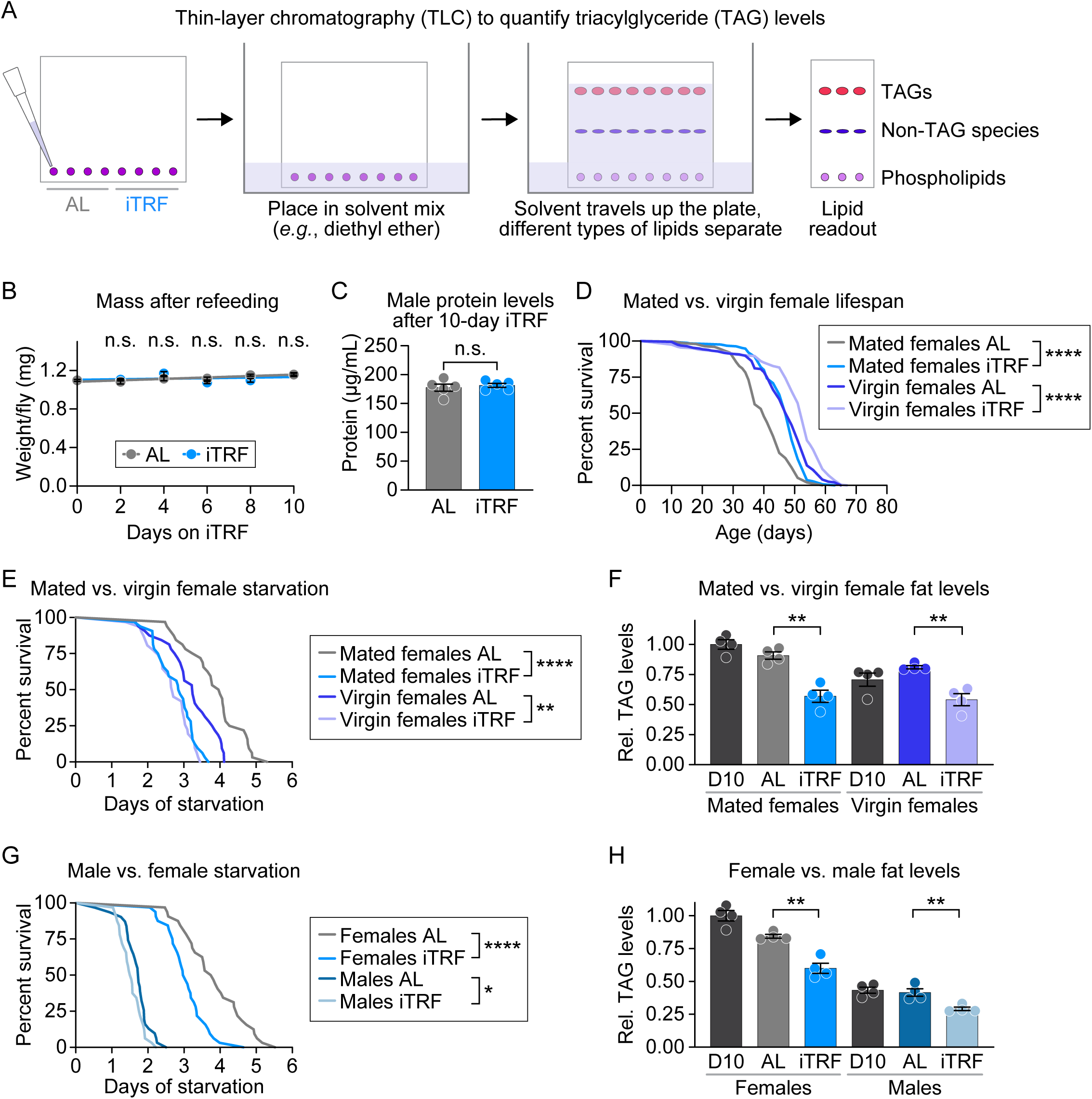
iTRF fat loss is not specific to sex or reproductive status. **(A)** Schematic of thin-layer chromatography (TLC) protocol to quantify triacylglyceride (TAG) levels. Samples are pipetted onto a silica-coated plate and placed in a solvent mix, which travels up the plate and separates different types of lipids, which can be quantified. **(B)** The mass of the flies remains unchanged throughout the 10-day iTRF period. Mass is measured as wet weight. n=10 flies/replicates/condition, with 4 replicates/condition; each point represents the average across replicates. p>0.05 compared to AL control for all iTRF time points measured. **(C)** iTRF does not affect protein levels in males. n=10 flies/replicate/condition, 5 replicates/ condition. p>0.05 compared to AL control. **(D)** iTRF extends the lifespan of both mated and virgin females. n=196-250 flies/condition, p<0.0001 compared to AL control for both virgin and mated females on iTRF. **(E)** Both mated and virgin females die faster during starvation on iTRF. n=32 flies/condition, p<0.01 compared to AL control for virgin females on iTRF, p<0.0001 compared to AL control for mated females on iTRF. **(F)** Both mated and virgin females lose fat on iTRF. n=10 flies/replicate/condition, 4 replicates/condition. p<0.01 compared to AL control for both mated and virgin females on iTRF. **(G)** Both males and females die faster during starvation on iTRF, n=32 flies/condition, p<0.0001 compared to AL control for females on iTRF, p<0.05 compared to AL control for males on iTRF. **(H)** Both males and females lose fat on iTRF. n=10 flies/replicate/condition for females, n=20 flies/replicate/condition for males, 4 replicates/condition, p<0.01 compared to AL control for both males and females on iTRF. p-values were obtained using independent t-tests unless otherwise indicated. Each experiment was performed 3 times.

**Supplemental Figure 2:**
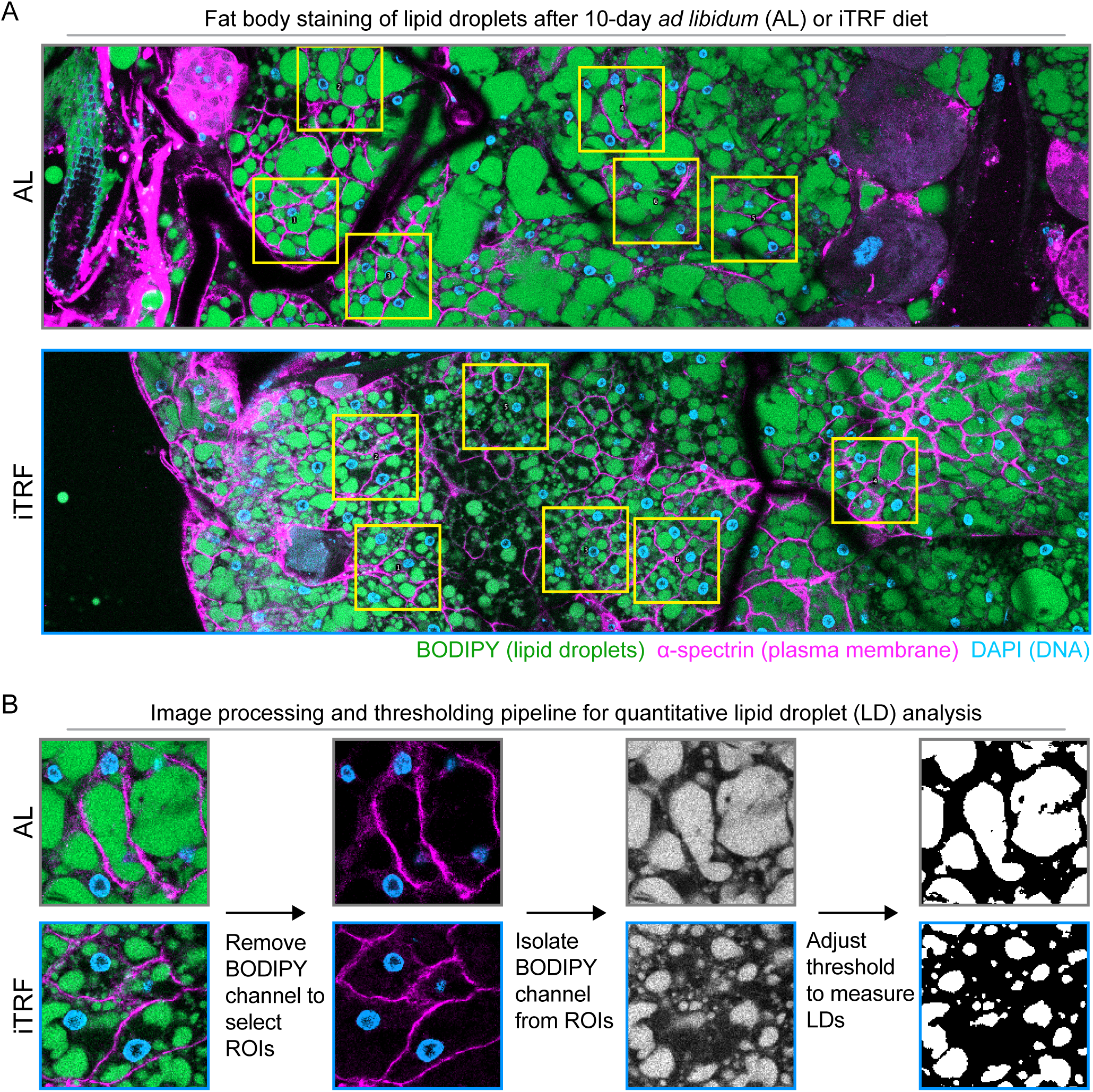
Fat body staining pipeline for quantitative lipid droplet analysis. **(A)** Representative images of fat body staining after 10-day AL or iTRF diets. green=BODIPY stain (fat), blue= DAPI stain (nuclei), magenta= ɑ-spectrin (membrane). **(B)** Image processing and thresholding pipeline for quantitative lipid droplet analysis. ROIs are chosen after the fat (BODPIY) channel is removed, using nuclei (DAPI) and plasma membranes (ɑ-spectrin) channels. After ROIs are chosen, the fat (BODPIY) channel is isolated, where thresholding is then applied to measure LD size.

**Supplemental Figure 3:**
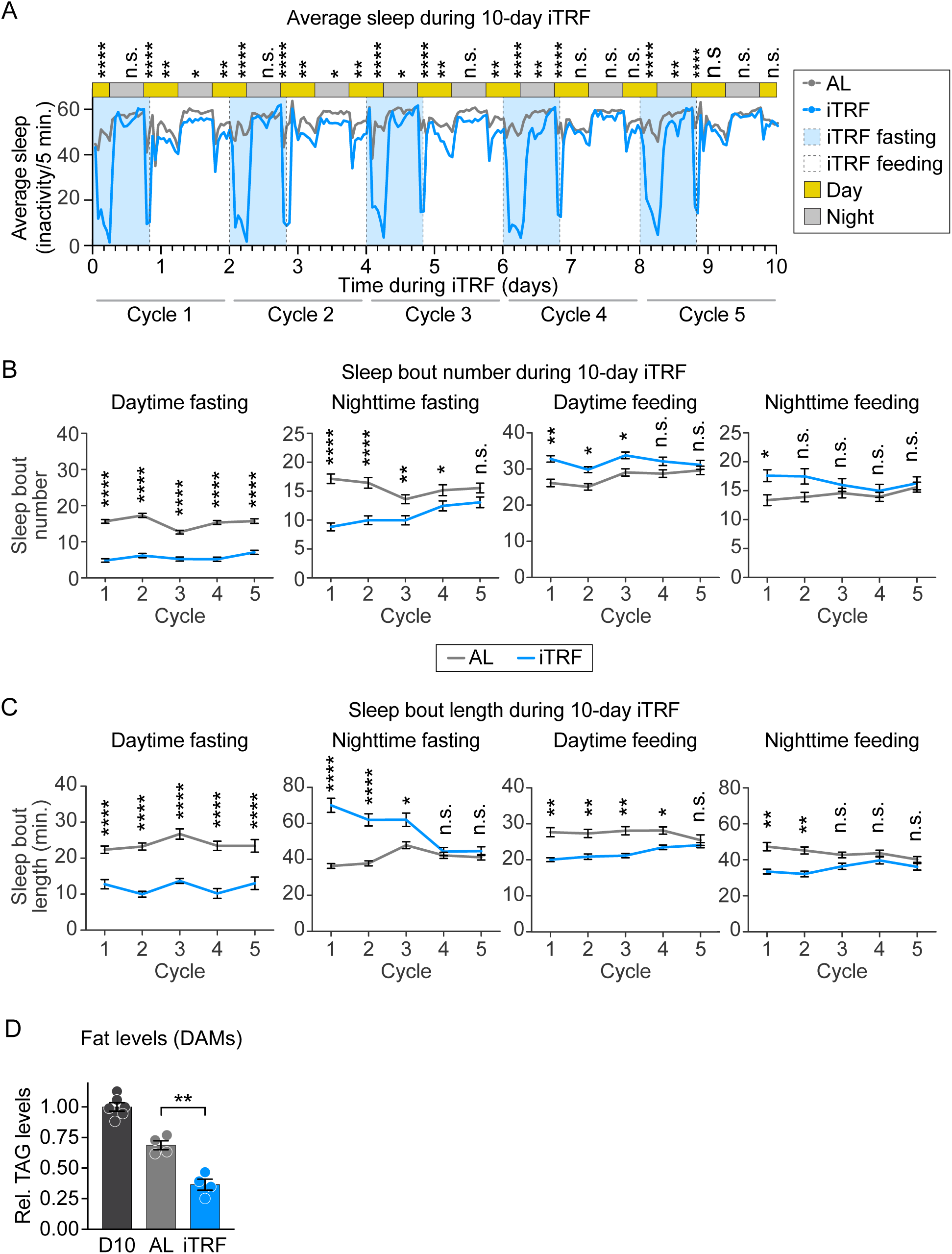
iTRF changes sleep patterns during the 10-day iTRF period. **(A)** Average sleep via DAM measurement with AL (grey) and iTRF (blue) plotted over the 10-day iTRF period. iTRF fasting periods (light blue shadow box) and feeding periods (white box) with day (yellow) and night (dark grey) boxes are also plotted. Lines represent flipping time for both AL and iTRF flies. p-values are generated by measuring the sleep of all flies under a condition, given the time of day and fasting/feeding state for iTRF flies. n=32 flies/replicate/ condition, with 3 replicates/condition for both AL and iTRF. **(B)** iTRF flies have reduced sleep bouts during day fasting and feeding. n=32 flies/replicate/condition with 3 replicates/ condition. **(C)** iTRF has shorter bout length during day fasting, but longer bout length in night fasting, day feeding, and night feeding toward the beginning of iTRF, which becomes non-significant in later cycles. n=32 flies/ replicate/condition, with 2-3 replicates/condition. p-values were obtained through independent t-tests unless otherwise noted. Each experiment was completed 2-3 times. **(D)** Confirmation of fat loss during iTRF DAM activity experiments. n=10 flies/replicate/condition, with 4 replicates/condition. p<0.01 compared to AL control for iTRF.

**Supplemental Figure 4:**
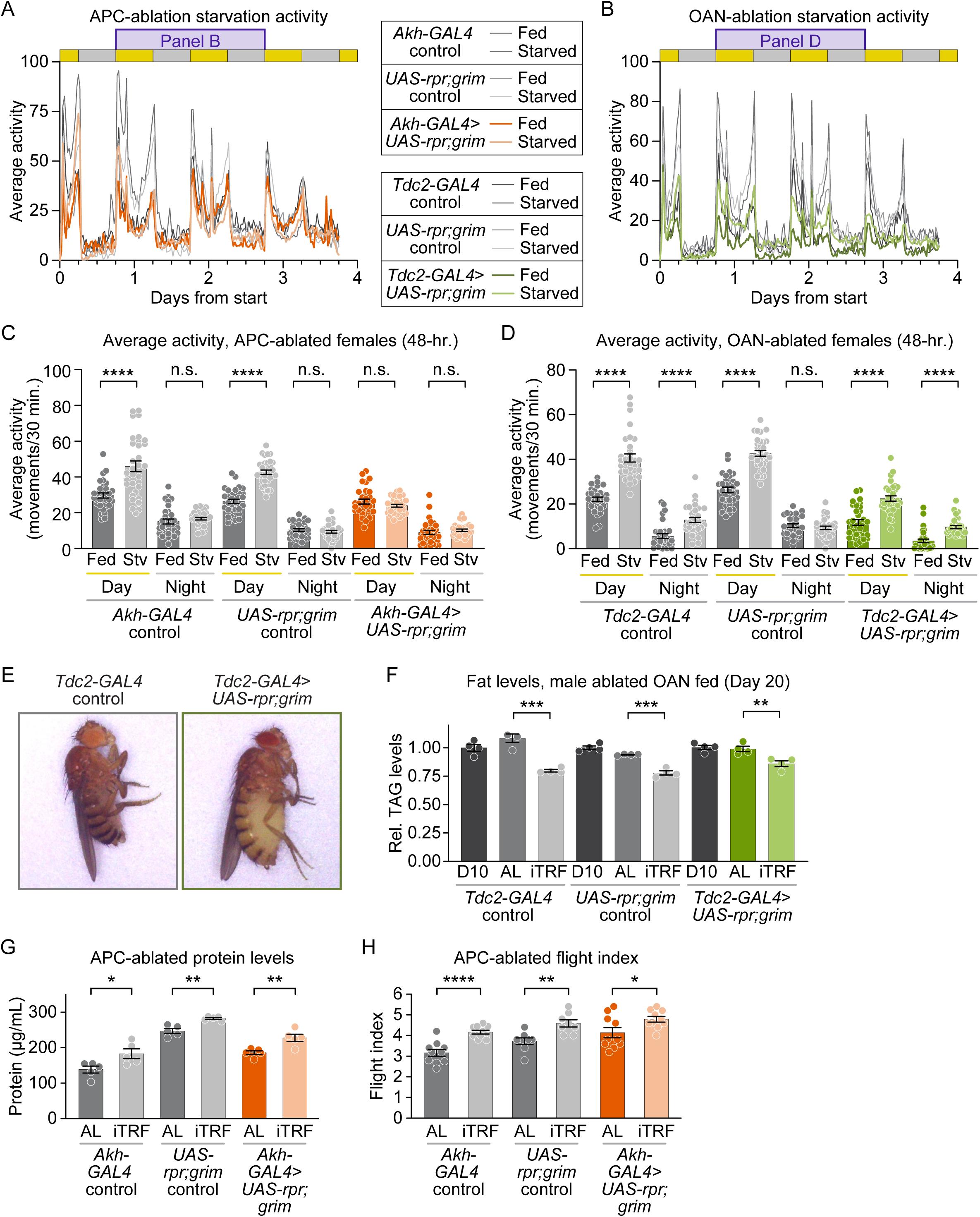
APC-and OAN-ablation leads to different physiological effects. **(A)** Starved and fed activity profiles of APC-ablated flies and their related controls. Day (yellow) and night (grey) boxes are noted on the graphs. **(B)** Starved and fed activity profiles of OAN-ablated flies and their related controls. Day (yellow) and night (grey) boxes are noted on the graphs. **(C)** Average activity for APC-ablated and genetic controls (starved and fed) during the denoted 48-hour period in **A**, as measured by day and night activity. **(D)** Average activity for OAN-ablated and genetic controls (starved and fed) during the 48-hour period in **B**, as measured by day and night activity. **(E)** *Tdc2-GAL4* control and *Tdc2-GAL4*> *UAS-rpr; grim* OAN-ablated females’ side-by-side, showing the extended abdomen of OAN-ablated females. **(F)** OAN-ablated males lose fat on iTRF, unlike females. 4 replicates/condition. p<0.001 compared to AL control for *Tdc2-GAL4* control on iTRF, p<0.001 compared to AL control for *UAS-rpr; grim* control on iTRF, p<0.01 compared to AL control for *Tdc2-GAL4*> *UAS-rpr; grim* OAN-ablated males on iTRF. **(G)** Like their genetic controls, APC-ablated flies gain protein on iTRF. n=10 flies/replicate/condition/genotype, with 5 replicates/condition/genotype. **(H)** Like their genetic controls, APC-ablated flies perform better on iTRF diet than AL. n=10 flies/replicate/ condition/genotype, with 8-10 replicates/condition/genotype.

